# HPV16 E6 oncoprotein promotes microhomology-mediated viral integration by upregulating POLQ expression

**DOI:** 10.1101/2025.11.20.689626

**Authors:** Guangli Zhu, Shuhei Asada, Jithma P Abeykoon, Lifang Sun, Sirisha Mukkavalli, Alan D. D’Andrea

## Abstract

Integration of the high-risk Human Papillomavirus 16 (HPV16) genome into the host chromosome, frequently driven by microhomology-mediated end joining (MMEJ), is a critical step in the carcinogenesis of HPV-associated tumors. However, the mechanisms by which viral oncoproteins manipulate the error-prone MMEJ pathway remain poorly defined. Here, we demonstrate that the HPV16 E6 oncoprotein upregulates MMEJ to facilitate viral genome integration. This heightened MMEJ activity is driven by a marked increase in the protein levels of DNA Polymerase Theta (POLQ), a central enzyme of the MMEJ pathway. Mechanistically, we show that the elevation of POLQ levels in response to HPV16 E6 expression is dependent on the host E3 ubiquitin ligase UBE3A/E6AP but is independent of p53 degradation. E6 activates UBE3A to promote the ubiquitination and degradation of RAD23A, a shuttle protein required for delivering polyubiquitinated POLQ to the proteasome. Consequently, the loss of functional RAD23A phenocopies the effect of HPV16 E6, leading to POLQ stabilization and increased MMEJ activity. By elucidating the E6-UBE3A-RAD23A-POLQ axis, our findings reveal a novel mechanism through which HPV manipulates the host DNA repair machinery to promote its integration and oncogenic potential.

## Introduction

Viral infections account for a significant proportion of human cancer cases, with human papillomavirus (HPV) being one of the most prevalent oncogenic viruses, particularly high-risk HPV types such as HPV16 (1). Globally, approximately 5% of all cancers, equivalent to 630,000 new cases annually, are attributed to HPV infection (2). HPV is the primary driver of cervical cancer (>95% of cases) and contributes to a wide range of other malignancies, including 88% of anal, 31% of oropharyngeal, 78% of vaginal, 25% of vulvar, and 50% of penile cancers, underscoring its broad oncogenic impact. Consequently, HPV-associated cancers represent a major global health burden.

HPV16 is one of the most oncogenic high-risk HPV viruses, driving malignant transformation primarily through the sustained expression of its oncoproteins E6 and E7 (3). HPV16 is a circular double-stranded DNA virus. The integration of the HPV genome into host DNA is a pivotal event in malignant transformation, ensuring the transition from transient infection to persistent infection. This transition enables the dysregulated expression of E6 and E7, which in turn disrupts cell cycle checkpoints to promote oncogenesis (4). However, despite its biological and clinical significance, the molecular pathways that mediate HPV integration remain poorly understood, particularly the role of host DNA repair pathways in this process.

For a circular double-stranded DNA virus like HPV to integrate into the host genome, two key prerequisites must be met: the presence of DNA double-strand breaks (DSBs) in both the viral and host genomes, and the engagement of DNA repair machinery. DSBs provide the necessary entry points for viral DNA insertion, and in HPV-associated cancers, these breaks frequently arise due to pre-existing genomic instability, particularly at fragile sites, actively transcribed regions, and areas of replication stress (5). Such regions are inherently prone to DNA breakage, making them preferential hotspots for viral integration.

Since HPV does not encode any DNA damage repair proteins, integration must rely entirely on the host cell’s DNA repair machinery to ligate the viral DNA with the broken host DNA. Mammalian cells employ multiple DNA repair pathways to resolve DSBs, including non-homologous end joining (NHEJ), homologous recombination (HR), and microhomology-mediated end joining (MMEJ) (6). While NHEJ is the predominant double-strand break (DSB) repair mechanism in mammalian cells, MMEJ has emerged as an alternative, error-prone pathway that utilizes 2-20 bp microhomologous sequences to mediate end joining (7). This pathway is dependent on the DNA polymerase theta (POLQ), encoded by *POLQ*. High-throughput sequencing of HPV-associated tumors has demonstrated that viral-host DNA junctions frequently contain microhomologous sequences. Microhomologous sequences flanking HPV integration sites exhibit distribution patterns characteristic of MMEJ-mediated repair, with most microhomologous regions localized around the integration center (8). Over 60% of HPV integration events may be mediated by MMEJ, reinforcing the idea that HPV preferentially integrates into DSBs repaired via this pathway (9). These findings suggest that HPV integration is not a random event but rather a process determined by the repair mechanisms active at the time of DNA breakage, with MMEJ playing a dominant role. We hypothesized that HPV16 E6 may enhance MMEJ activity by modulating key host regulatory factors, thereby facilitating viral genome integration.

In the current study, we demonstrate that HPV16 E6 strongly enhances MMEJ activity and facilitates viral DNA integration, an effect significantly more pronounced than that of E7. Mechanistically, we show that HPV16 E6 elevates POLQ protein level and promotes MMEJ repair in a manner dependent on UBE3A (also known as E6AP), a key E3 ubiquitin ligase activated by E6 (10), but independent of p53. HPV16 E6-enhanced ubiquitination and degradation of RAD23A by UBE3A prevents RAD23A, a proteasomal shuttle protein (11, 12), from delivering ubiquitinated POLQ to the proteasome. Reduced proteosome delivery results in increased POLQ protein levels and increased MMEJ activity. By elucidating the role of HPV16 E6 in modulating MMEJ, our findings reveal a novel mechanism through which HPV manipulates host DNA repair machinery and provide new insights into the molecular basis of HPV genome integration.

## Results

### HPV16 E6 overexpression enhances MMEJ activity and MMEJ-mediated double-stranded DNA integration efficiency

To investigate the impact of HPV16 E6 overexpression on MMEJ pathway, we achieved stable expression of HA-tagged HPV16 E6 or E7 via lentiviral transduction in U2OS cells (**Fig. 1A**). Using a previously reported ISce-I endonuclease-based MMEJ-GFP reporter system (13), we found that HPV16 E6 expression caused a marked increase in MMEJ activity, whereas HPV16 E7 expression exhibited only modest effect (**Fig. 1B**). Notably, HPV16 E6 ΔN6, an alternative translation product of E6 that lacks six N-terminal residues (14), further enhanced MMEJ activity compared to full-length E6 (**Fig. S1A**). In contrast, neither E6 nor E7 affected NHEJ efficiency (**Fig. S1B**), and E6 induced only a slight increase in HR activity (**Fig. S1C**).

**Figure 1.**
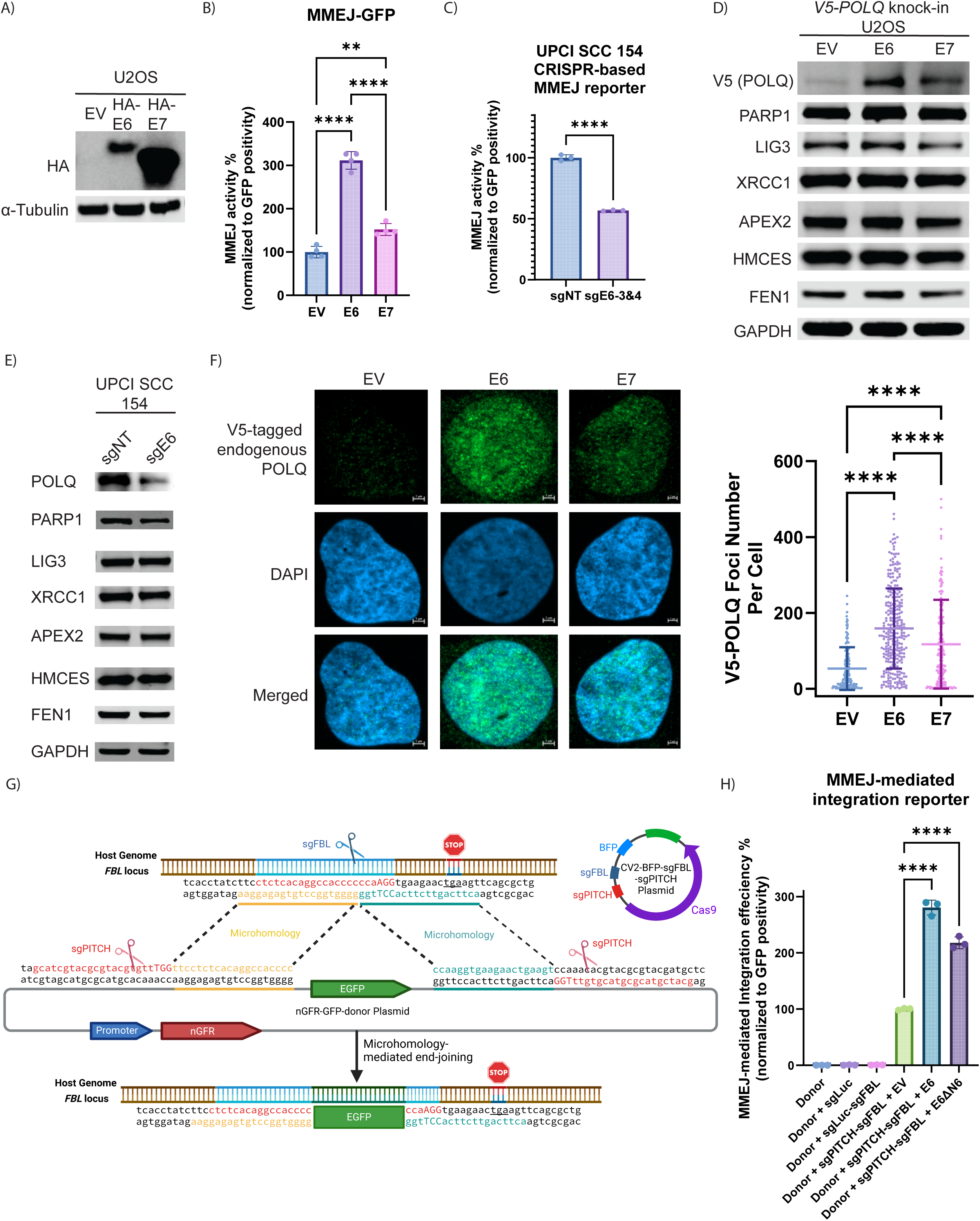
HPV16 E6 enhances MMEJ pathway activity and MMEJ-mediated DNA integration efficiency. A) Immunoblot of HA in U2OS cells expressing empty vector (EV), HA-tagged HPV16 E6, or E7. B) Relative microhomology-mediated end joining (MMEJ) activity measured using the I-SceI-based MMEJ reporter assay in U2OS cells expressing EV, HPV16 E6, or HPV16 E7. Data are shown as the mean ± SD. Statistical significance was calculated using one-way ANOVA with Tukey’s post hoc test (**, *P* < 0.01; ****, *P* < 0.0001). C) Relative MMEJ activity evaluated by a CRISPR-Cas9-based reporter assay in HPV-positive UPCI-SCC-154 cells with or without HPV16 *E6* knock-out. Data are shown as the mean ± SD. Statistical significance was calculated using two-tailed Student’s t-test (****, *P* < 0.0001). D) Immunoblot of V5-tagged endogenous POLQ, other key MMEJ pathway components (PARP1, LIG3, XRCC1, APEX2, HMCES, FEN1) and GAPDH in *V5*-*POLQ* knock-in U2OS cells expressing empty vector, HPV16 E6, or HPV16 E7. E) Immunoblot of endogenous POLQ, other MMEJ pathway components and GAPDH in HPV-positive UPCI-SCC-154 cells with or without HPV16 *E6* knock-out. F) Immunofluorescence analysis showing representative images (Scale bar = 2 μm) and quantification of V5-tagged endogenous POLQ and γ-H2AX foci formation in V5-*POLQ* knock-in U2OS cells expressing empty vector, HPV16 E6, or HPV16 E7. At least 150 cells were counted. Data are shown as the mean ± SD. Statistical significance was calculated using one-way ANOVA with Tukey’s post hoc test (****, *P* < 0.0001). G) Schematic representation of the reporter system designed to measure MMEJ-mediated double-strand DNA integration efficiency. H) Relative MMEJ-mediated integration efficiency in HEK293T cells expressing HPV16 E6 or control vector. Controls included cells transfected with donor DNA alone, donor DNA plus non-targeting sgRNA (sgLuc), or donor DNA plus both sgLuc and sgFBL. Data are shown as the mean ± SD. Statistical significance was calculated using two-tailed Student’s t-test (****, *P* < 0.0001).

To further validate the effect of HPV16 E6 on MMEJ activity, we developed a CRISPR-based MMEJ reporter system (**Fig. S1D**). *POLQ* knock-out or pharmacological inhibition of POLQ using ART558 significantly reduced MMEJ activity, confirming the specificity of this assay to measure POLQ-dependent MMEJ (**Fig. S1E**). As measured by this CRISPR-based MMEJ reporter system, E6 expression significantly increased MMEJ activity in the HPV-negative FaDu cells (**Fig. S1F**), and depletion of E6 in HPV16-positive UPCI-SCC-154 cells significantly reduced MMEJ activity (**Fig. 1C**), confirming that endogenous E6 contributes to MMEJ activation in HPV-positive cells.

To determine which MMEJ factor is most likely responsible for the E6-mediated MMEJ activation, we profiled the protein expression of key MMEJ factors (7). Immunoblotting of *V5*-*POLQ* knock-in U2OS cells revealed that HPV16 E6 specifically increased POLQ protein levels, while other MMEJ-related proteins (APEX2, HMCES, FEN1, LIG3, PARP1, XRCC1) remained unchanged (**Fig. 1D**). These effects were also observed in the HPV-negative HNSCC cell lines-CAL33 and SCC9 following ectopic expression of E6 (**Fig. S1G-H**). Conversely, *E6* knock-out in HPV-positive UPCI-SCC-154 cells reduced POLQ protein level (**Fig. 1E**). Immunofluorescence further demonstrated a significant increase in POLQ nuclear foci assembly in E6-expressing cells (**Fig. 1F**). However, HPV16 E6 expression did not significantly alter *POLQ* mRNA levels (**Fig. S1I-J**), indicating that the E6 increases POLQ protein level through post-transcriptional mechanisms.

Next, we investigated whether E6 expression enhances the efficiency of MMEJ-mediated double-stranded DNA integration. Since it is difficult to generate live HPV virus *in vitro*, we established an MMEJ-mediated double-stranded DNA integration reporter based on the PITCH MMEJ-mediated knock-in system (15) (**Fig. 1G**). This reporter assay demonstrated that both E6 and E6 DN6 significantly increased the MMEJ-mediated integration efficiency of donor DNA fragments into the host genome (**Fig. 1H**).

Taken together, these results show that HPV16 E6 enhances MMEJ repair activity, selectively elevates POLQ protein level, and promotes MMEJ-mediated double-stranded DNA integration efficiency, suggesting a key role for E6 in facilitating HPV genome integration through the MMEJ pathway.

### Hyperactivation of MMEJ driven by the HPV16 E6 oncoprotein overcomes mitotic DNA damage caused by increased replication stress

Since DSBs are a prerequisite for viral integration, we next investigated the impact of HPV16 E6 on genomic stability. We assessed DNA damage levels using γ-H2AX as a marker of DSBs and phosphorylated replication protein A (pRPA) as an indicator of replication stress. HPV16 E6 expression significantly increased both γ-H2AX and pRPA levels, with a more pronounced effect than HPV16 E7 (**Fig. 2B**). These findings were corroborated by the alkaline comet assay, which revealed increased tail length in E6-expressing cells, indicative of elevated DNA fragmentation (**Fig. 2A**).

**Figure 2.**
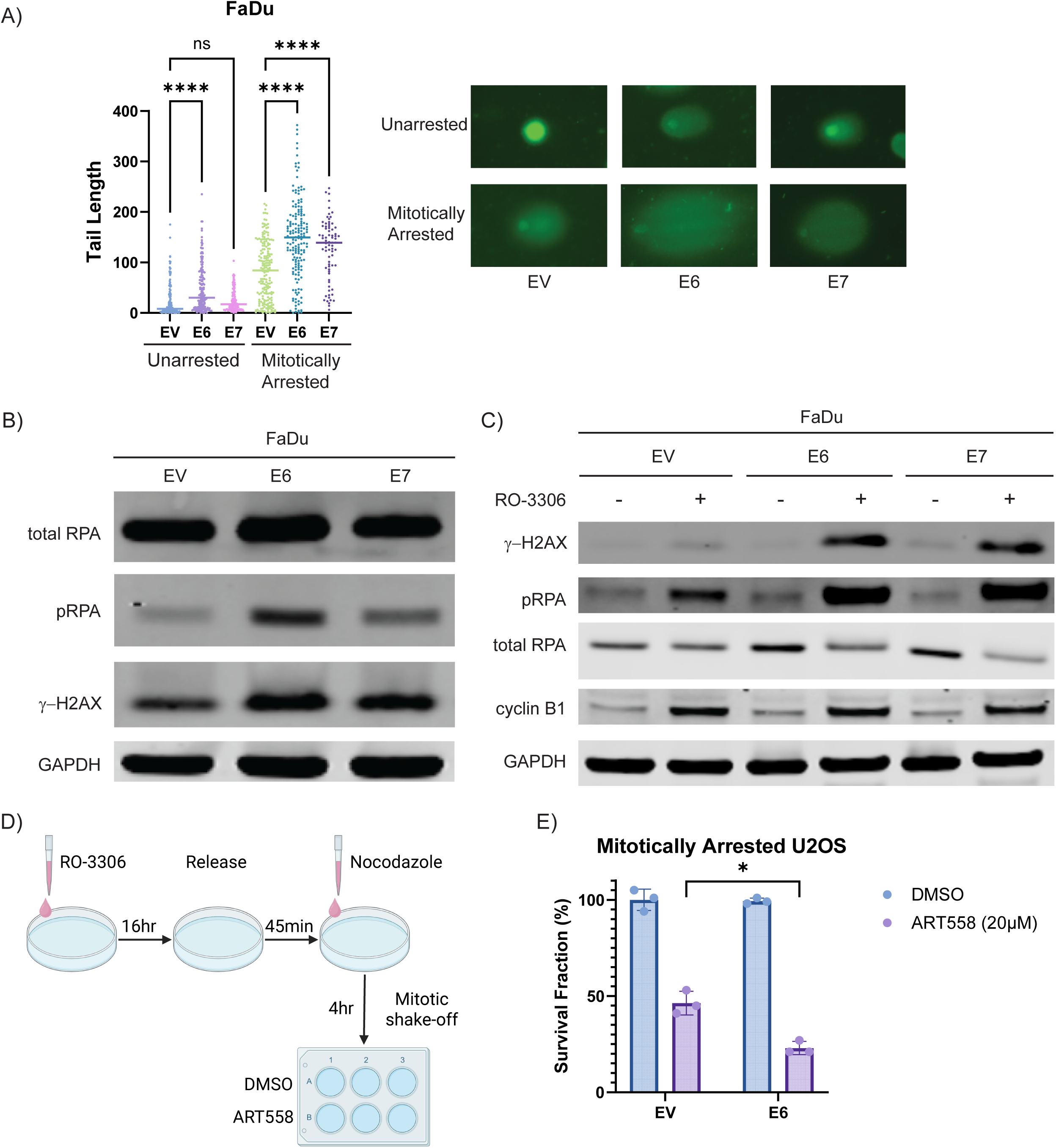
HPV16 E6 induces DNA damage and replication stress, particularly during mitosis. A) Quantification of DNA double-strand breaks using alkaline comet assay in asynchronous or mitotically arrested FaDu cells expressing empty vector, HPV16 E6, or HPV16 E7. Data are shown as the mean ± SD. Statistical significance was calculated using two-tailed Student’s t-test (*, *P* < 0.05). B) Immunoblot of replication stress markers (phospho-RPA S33, total RPA), DNA damage marker (γ-H2AX), and GAPDH in FaDu cells expressing empty vector, HPV16 E6, or HPV16 E7. C) Immunoblot of DNA damage response proteins (phospho-RPA S33, total RPA, γ-H2AX), mitotic marker (cyclin B1), and GAPDH in asynchronous or mitotically arrested FaDu cells expressing empty vector, HPV16 E6, or HPV16 E7. D) Schematic illustration of clonogenic assay for mitotically arrested cells. E) Clonogenic survival assay of mitotically arrested U2OS cells expressing empty vector or HPV16 E6, treated continuously with DMSO or 20 µM ART558. Data are shown as the mean ± SD. Statistical significance was calculated using one-way ANOVA with Tukey’s post hoc test (****, *P* < 0.0001).

To determine whether the increased replication stress in cells expressing the E6 oncoprotein leads to mitotic entry with unrepaired DSBs, FaDu cells were synchronized and arrested in M phase using RO-3306 and nocodazole. Mitotic arrest was confirmed by elevated levels of the mitotic marker, cyclin B1. Following synchronization, HPV16 E6-expressing cells exhibited further elevations in γ-H2AX and pRPA levels, suggesting that E6-induced DNA damage accumulates during mitosis (**Fig. 2C**). This was further supported by the comet assay, which revealed a significant increase of DNA fragmentation in mitotic cells expressing E6 compared to controls (**Fig. 2A**). Given that MMEJ is the dominant DSB repair pathway during mitosis (16–18), we next asked whether E6-expressing cells depend on POLQ for survival under these conditions. Indeed, clonogenic survival assays demonstrated that mitotically arrested E6-expressing cells were markedly more sensitive to the POLQ inhibitor ART558 than control cells (**Fig. 2D-E**), suggesting a heightened dependency on the MMEJ pathway for survival under these conditions.

These observations support a model in which HPV16 E6-induced replication stress leads to the accumulation of DSBs, particularly during mitosis, which may contribute to its role in promoting genomic instability and facilitating viral genome integration. Recent publications have shown that MMEJ is the primary active DSB repair pathway during mitosis, further supporting a link between HPV16 E6-induced DNA damage accumulation during mitosis and the cellular reliance of MMEJ for DSB repair (16–18).

### Upregulation of MMEJ activity by HPV16 E6 is dependent on UBE3A

We next investigated the underlying mechanisms of the HPV16 E6-induced MMEJ activation. Since HPV16 E6 promotes degradation of p53 by acting as an allosteric activator of UBE3A (10), we first tested whether E6-mediated MMEJ activation is dependent on p53 suppression. To this end, we knocked out *TP53* in U2OS cells and measured MMEJ activity, using the MMEJ-GFP reporter assay in the presence or absence of E6 overexpression (**Fig. 3A**). While *TP53* knock-out alone increased MMEJ activity, which is consistent with previous studies (19), E6 overexpression further enhanced the MMEJ activity in *TP53* knock-out cells, indicating that E6 promotes MMEJ through a mechanism independent of p53 degradation (**Fig. 3B**). Similarly, the elevated MMEJ activity in wild-type HPV-positive UPCI-SCC-154 cells was not abolished by *TP53* knock-out (**Fig. 3D**).

**Figure 3.**
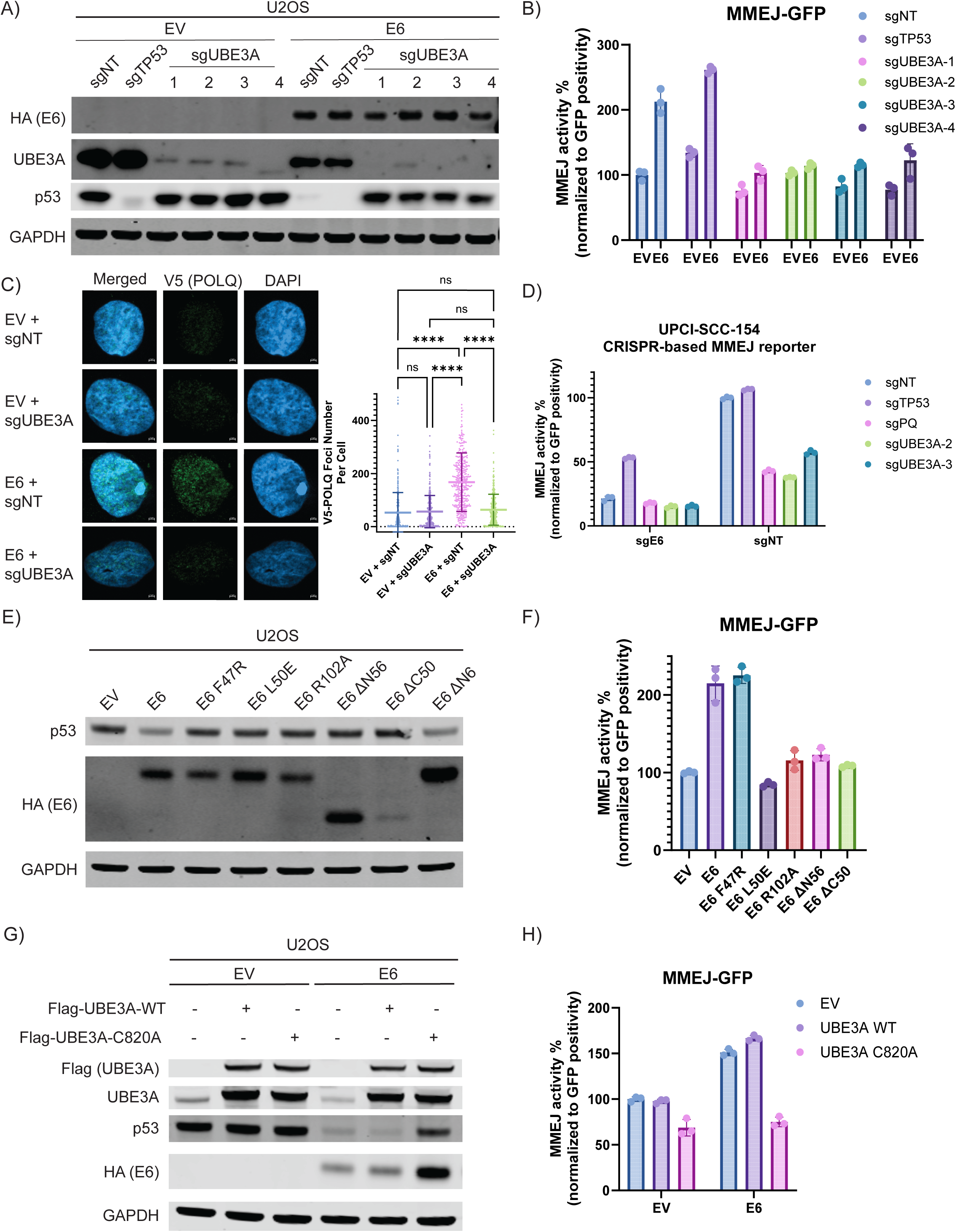
HPV16 E6 promotes MMEJ repair through a UBE3A-dependent and p53-independent mechanism A) Immunoblot of UBE3A, p53, HA-tagged HPV16 E6, and GAPDH proteins in wild-type, *TP53* knock-out, or *UBE3A* knock-out U2OS cells expressing empty vector or wild-type HPV16 E6. B) Relative microhomology-mediated end joining (MMEJ) activity measured using the I-SceI-based MMEJ GFP reporter assay in wild-type, *TP53* knock-out, or *UBE3A* knock-out U2OS cells expressing empty vector or wild-type HPV16 E6. C) Immunofluorescence analysis showing representative images and quantification of V5-tagged endogenous POLQ and γ-H2AX foci formation in wild-type or UBE3A knock-out V5-*POLQ* knock-in U2OS cells expressing empty vector, HPV16 E6, or HPV16 E7. At least 150 cells were counted. Data are shown as the mean ± SD. Statistical significance was calculated using one-way ANOVA with Tukey’s post hoc test (****, *P* < 0.0001). D) Relative MMEJ activity evaluated by CRISPR-Cas9-based reporter assay in wild-type, *TP53* knock-out, or *UBE3A* knock-out HPV-positive UPCI-SCC-154 cells with or without HPV16 *E6* knock-out. E) Immunoblot of p53, HA-tagged wild-type and mutant HPV16 E6, and GAPDH proteins in MMEJ reporter U2OS cells expressing the indicated E6 constructs. F) Relative MMEJ activity measured using the I-SceI-based MMEJ GFP reporter assay in U2OS cells expressing empty vector, wild-type or mutant HPV16 E6 proteins. G) Immunoblot of FLAG-tagged wild-type or catalytically inactive (C820A) UBE3A, p53, HA-tagged E6, and GAPDH in U2OS cells expressing empty vector, wild-type or C820A mutant UBE3A, with or without HPV16 E6 expression. H) Relative MMEJ activity measured using the I-SceI-based MMEJ GFP reporter assay in U2OS cells expressing empty vector, wild-type UBE3A or C820A mutant, with or without HPV16 E6 expression.

To determine whether UBE3A is required for E6-induced MMEJ activation, we next performed *UBE3A* knock-out in U2OS cells. Strikingly, the deletion of UBE3A abolished the E6-induced enhancement of MMEJ activity, suggesting that UBE3A is essential for this process (**Fig. 3B**). Consistent with the effect on MMEJ activity, *UBE3A* knock-out also decreased POLQ foci assembly and the POLQ protein level in E6-overexpressing cells (**Fig. 3C, Fig. S3A**). Similar results were observed in HPV16-positive UPCI-SCC-154 cells, where the MMEJ activation was abrogated by *UBE3A* knock-out (**Fig. 3D**). A comparable dependency on UBE3A was also observed in the MMEJ-mediated integration reporter assay, confirming that UBE3A, rather than loss of p53, is responsible for E6-mediated activation of MMEJ (**Fig. S2A-B**).

To further dissect the molecular basis of this regulation, we examined a series of E6 mutants with altered binding affinities to UBE3A or p53 (20, 21), which was validated by rescue of p53 expression (**Fig. 3E**). As expected, the E6 ΔN56 and E6 ΔC50 mutants, which are defective in both UBE3A and p53 binding, failed to enhance MMEJ activity (**Fig. 3F**). In contrast, the E6 F47R mutant, which retains UBE3A interaction but loses p53 binding, was still able to enhance MMEJ activity to a similar extent as wild-type E6, supporting the idea that p53 suppression is dispensable for MMEJ activation (**Fig. 3F**). Importantly, the E6 mutants specifically defective in UBE3A binding, including E6 L50E and E6 R102A, failed to enhance MMEJ activity, demonstrating that the interaction between E6 and UBE3A is necessary for E6’s effect on MMEJ pathway (**Fig. 3F**). Similar effects were also observed in the mutants of an alternative translation product of E6, E6 ΔN6 (**Fig. S3B-D**). Consistently, the same effects were also detected using the MMEJ-mediated integration reporter (**Fig. S2C**).

To further investigate whether the E3 ligase activity of UBE3A is required for E6-mediated MMEJ activation, we mutated the catalytic cysteine residue of UBE3A to alanine (C820A), which disrupts the catalytic activity of UBE3A while preserving its ability to interact with substrates. We reintroduced either wild-type or C820A UBE3A into U2OS cells and measured MMEJ activity in the presence of E6 overexpression. The dominant-negative effect of UBE3A-C820A was evident, as indicated by the recovery of p53 levels in cells co-expressing E6 and UBE3A-C820A (**Fig. 3G**). The UBE3A-C820A mutant abolished E6-induced MMEJ enhancement, indicating that the catalytic activity of UBE3A is necessary for E6-mediated MMEJ enhancement (**Fig. 3H**). A similar requirement for the catalytic activity of UBE3A was observed in the MMEJ-mediated integration assay (**Fig. S2D**). Again, the alternative translation product E6 ΔN6 also exhibited the same phenotype as the full-length E6 (**Fig. S3E-F**).

Collectively, these findings establish that HPV16 E6 enhances MMEJ activity through a UBE3A-dependent but p53-independent mechanism. UBE3A is therefore a key regulator of MMEJ activity in HPV16-positive cells, revealing a previously unrecognized function of this E3 ligase in DNA repair pathway modulation.

### Ubiquitination of RAD23A by UBE3A mediates the impact of E6 on MMEJ

Having established that HPV16 E6 promotes MMEJ activity by activating UBE3A, we next sought to identify the specific UBE3A substrate responsible for this effect. A previous study has identified RAD23A, a ubiquitin-binding protein involved in nucleotide excision repair and proteasomal degradation, as a substrate of UBE3A (22–24). Furthermore, the ubiquitination of RAD23A by UBE3A is enhanced by E6 (22, 23, 25). Using co-immunoprecipitation, we demonstrated an enhanced interaction between UBE3A and RAD23A upon E6 expression (**Fig. 4A**), supporting a model in which E6 facilitates RAD23A ubiquitination through UBE3A activation.

**Figure 4.**
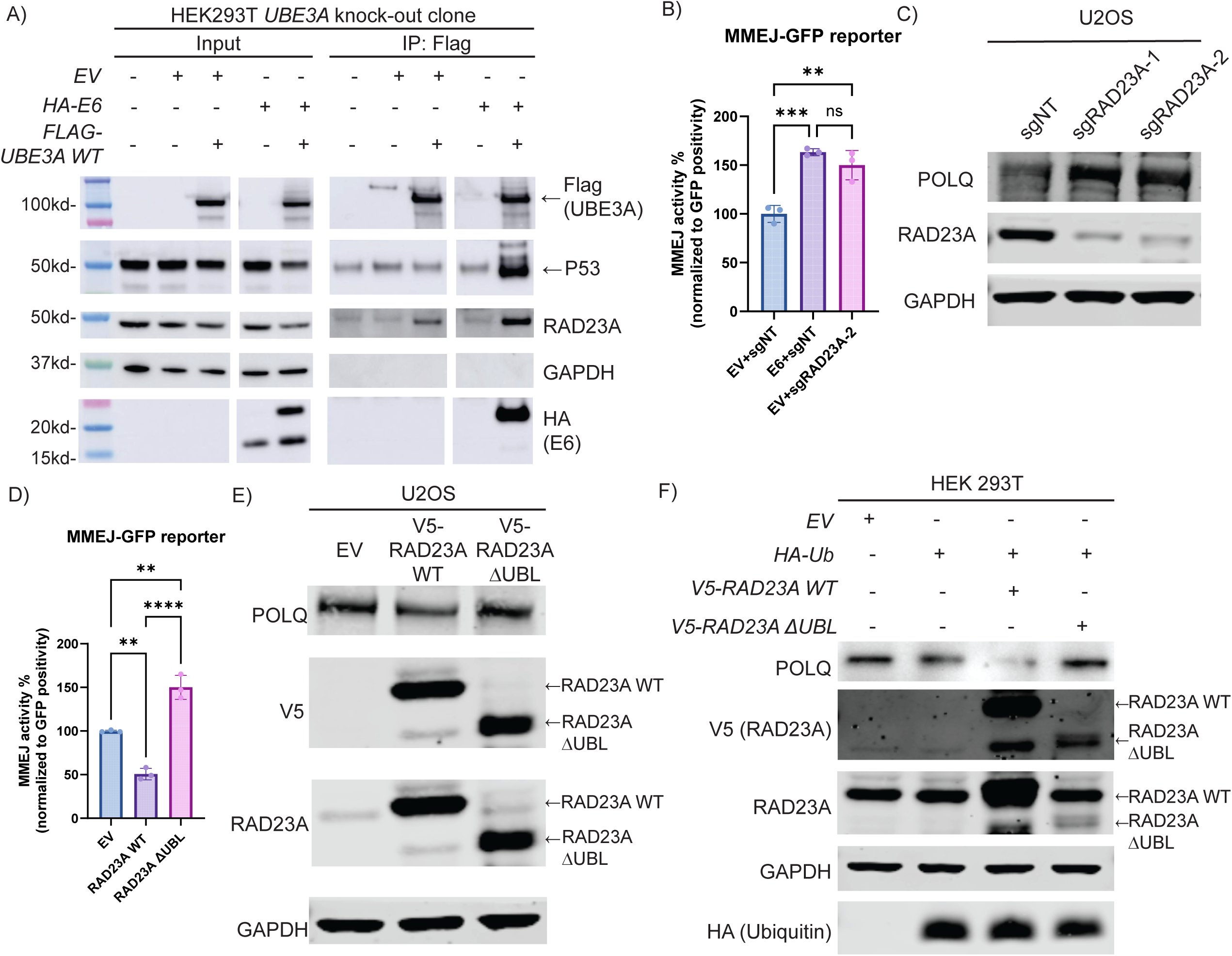
RAD23A mediates the impact of E6 on MMEJ activity by regulating POLQ stability A) Co-immunoprecipitation analysis. Whole cell lysates from HEK293T cells transduced with indicated constructs were immunoprecipitated (IP) with anti-FLAG antibody, followed by Immunoblot of input and IP samples for FLAG (UBE3A), HA (HPV16 E6), p53, and RAD23A. B) Relative microhomology-mediated end joining (MMEJ) activity measured using the I-SceI-based MMEJ GFP reporter assay in wild-type or *RAD23A* knock-out U2OS cells with or without E6 overexpression. Data are shown as the mean ± SD. Statistical significance was calculated using one-way ANOVA with Tukey’s post hoc test (**, *P* < 0.01; ***, *P* < 0.001). C) Immunoblot of endogenous POLQ, RAD23A, and GAPDH in U2OS cells with or without *RAD23A* knock-out. D) Relative MMEJ activity measured using the I-SceI-based MMEJ GFP reporter assay in U2OS cells overexpressing empty vector, wild-type RAD23A or RAD23A ΔUBL mutant. Data are shown as the mean ± SD. Statistical significance was calculated using one-way ANOVA with Tukey’s post hoc test (**, *P* < 0.01; ****, *P* < 0.0001). E) Immunoblot of endogenous POLQ, V5 (RAD23A), RAD23A, UBE3A and GAPDH in U2OS cells overexpressing empty vector, wild-type RAD23A or RAD23A ΔUBL mutant. F) Immunoblot of endogenous POLQ, V5 (RAD23A), RAD23A, ubiquitin and GAPDH in HEK293T cells overexpressing empty vector, and ubiquitin together with empty vector, wild-type RAD23A or RAD23A ΔUBL mutant.

To determine whether RAD23A is involved in E6-mediated MMEJ activation, we performed the MMEJ-GFP reporter assay to assess MMEJ activity in wild-type and *RAD23A* knock-out U2OS cells. Notably, *RAD23A* knock-out alone significantly increased MMEJ activity, phenocopying the effect of E6 overexpression (**Fig. 4B**). Given that HPV16 E6 elevates POLQ expression, we next investigated whether RAD23A also regulates POLQ levels. Immunoblot showed that the loss of RAD23A increased POLQ protein levels (**Fig. 4C**), suggesting that RAD23A functions as a negative regulator of POLQ stability. Furthermore, overexpression of wild-type RAD23A suppressed MMEJ activity, whereas the RAD23A ΔUBL mutant, which lacks the ubiquitin-like (UBL) domain required for proteasome interaction (26–28), failed to inhibit MMEJ and instead acted in a dominant-negative manner (**Fig. 4D**). Consistently, wild-type RAD23A overexpression decreased POLQ protein levels, while the ΔUBL mutant had little effect (**Fig. 4E**).

To further assess whether RAD23A facilitates POLQ degradation in a ubiquitin-dependent manner, we co-expressed ubiquitin together with either wild-type or ΔUBL RAD23A. While wild-type RAD23A further decreased POLQ levels in the presence of ubiquitin, the ΔUBL mutant not only failed to reduce POLQ abundance but also reversed the ubiquitin-induced suppression (**Fig. 4F**). Collectively, these results demonstrate that RAD23A mediates the ubiquitin-dependent degradation of POLQ.

The results above strongly suggested that POLQ itself is a substrate for ubiquitin-mediated proteasomal degradation. Supporting this, treatment with MG132, a proteasome inhibitor, markedly increased POLQ protein levels (**Fig. S4A**). To further assess whether POLQ undergoes direct ubiquitination, we performed co-immunoprecipitation of FLAG-tagged POLQ from HEK293T cells with or without co-expression of HA-tagged ubiquitin. Immunoblot revealed that POLQ undergoes both mono- and polyubiquitination. Inhibition of the proteasome with MG132 led to a marked accumulation of polyubiquitinated POLQ, consistent with the idea that ubiquitinated POLQ is normally destined for proteasomal degradation (**Fig. 5A**).

**Figure 5.**
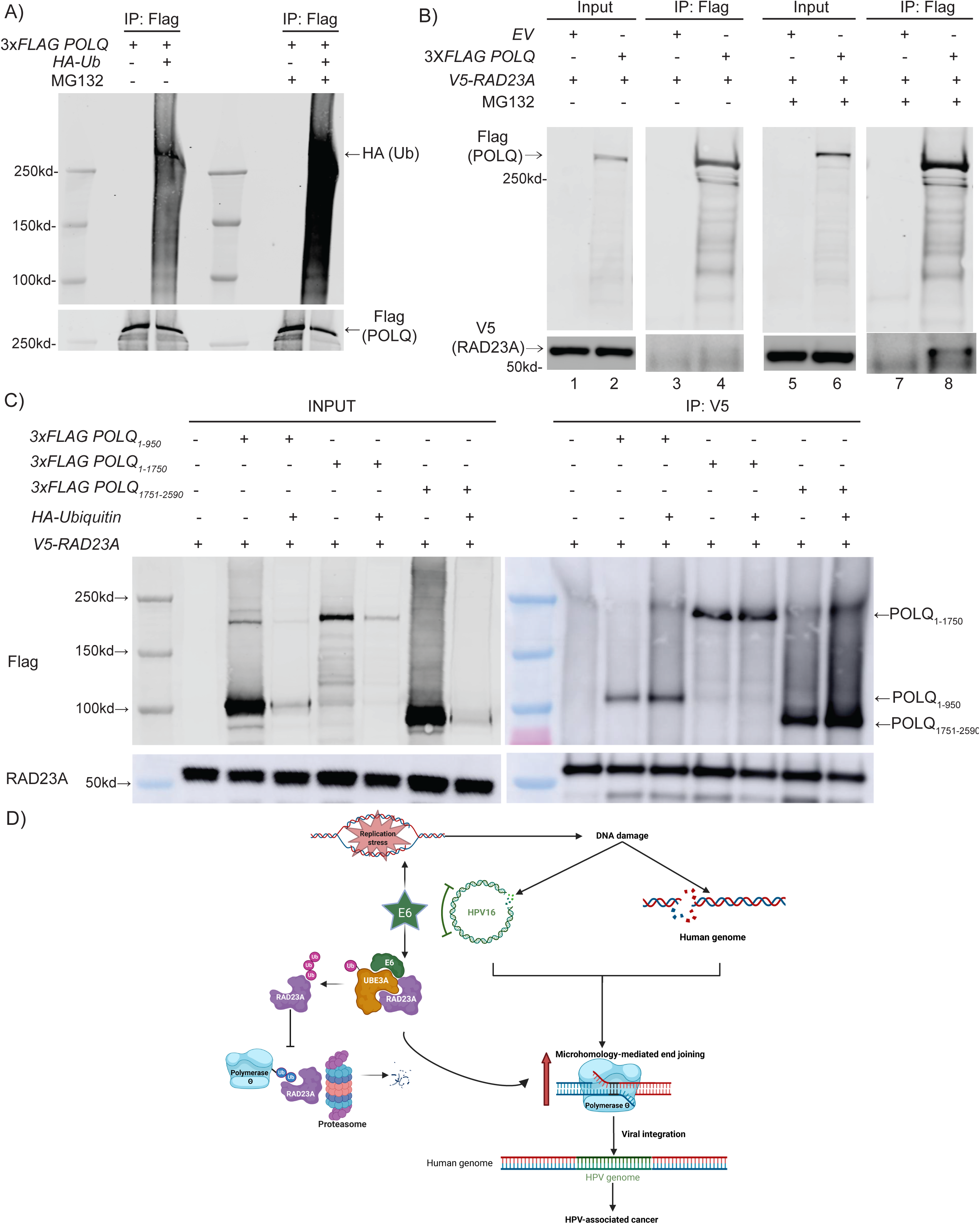
RAD23A interacts with POLQ and promotes its ubiquitination. A) Co-immunoprecipitation analysis. Whole cell lysates from POLQ knock-out HEK293T cells reconstituted with empty vector or HA-Ub, together with 3xFLAG-tagged full-length POLQ, with or without MG132 treatment, were immunoprecipitated with anti-Flag antibody, followed by immunoblotting of input and IP samples for Flag (POLQ) and HA (Ubiquitin). B) Co-immunoprecipitation analysis. Whole cell lysates from POLQ knock-out HEK293T cells reconstituted with empty vector or 3xFLAG-tagged full-length POLQ, together with V5-RAD23A, with or without MG132 treatment, were immunoprecipitated with anti-Flag antibody, followed by immunoblotting of input and IP samples for Flag (POLQ) and V5 (RAD23A). C) Co-immunoprecipitation analysis. Whole cell lysates from POLQ knock-out HEK293T cells reconstituted with empty vector, 3xFLAG-tagged full-length POLQ or its truncated mutants (1- 950, 1-1750, 1751-2590), together with V5-RAD23A, with or without HA-ubiquitin, were immunoprecipitated with anti-V5 antibody, followed by immunoblotting of input and IP samples for Flag (POLQ fragments) and RAD23A. D) Schematic of the working model for HPV16 E6-enhanced MMEJ-mediated viral integration.

Finally, to determine if RAD23A directly recognizes ubiquitinated POLQ, we expressed 3x*FLAG*-tagged full-length *POLQ* with or without *V5*-tagged *RAD23A* in a *POLQ* knock-out HEK293T clone. Co-immunoprecipitation with an anti-FLAG antibody revealed that, in the absence of MG132, only weak binding of V5-tagged RAD23A to full-length POLQ was observed (**Fig. 5B**). Upon MG132 treatment, however, the binding of V5-tagged RAD23A to full-length POLQ was markedly enhanced (**Fig. 5B, lane 8**), confirming that RAD23A preferentially recognizes ubiquitinated POLQ to mediate its proteasomal degradation. Similarly, the same phenomenon was recapitulated between endogenous RAD23A and POLQ (**Fig. S4B**). Reciprocal co-immunoprecipitation using an anti-V5 antibody further revealed that both the N-terminal and C-terminal regions of POLQ are subject to ubiquitination (**Fig. 5C**).

Taken together, these findings identify RAD23A as a key mediator of E6-induced MMEJ activation. Our results suggest that HPV16 E6 promotes error-prone DNA repair by facilitating the UBE3A-mediated ubiquitination of RAD23A, thereby impairing RAD23A-mediated degradation of POLQ and ultimately enhancing POLQ protein levels and MMEJ activity (**Fig. 5D**).

## Discussion

The integration of HPV DNA into the host genome is a pivotal event in the pathogenesis of HPV-associated cancers. However, the precise molecular mechanisms governing this process remain poorly understood. Our study shows that E6-induced DNA damage and MMEJ activation are key mechanisms exploited by HPV16 E6 to facilitate viral integration. A central finding of this investigation is the dominant role of HPV16 E6 in driving MMEJ activation, which supersedes the influence of E7, despite previous studies implicating E7 in the modulation of DNA repair pathways (29). We demonstrated that this process is mediated through a previously unrecognized mechanism involving UBE3A-dependent ubiquitination of RAD23A, a negative regulator of POLQ, thereby promoting MMEJ activity.

Accumulating evidence supports MMEJ as the predominant pathway exploited by HPV for genomic integration. Unlike retroviruses, HPV lacks an integrase and relies on host DNA repair machinery to mediate genome integration. A striking feature of HPV integration is the frequent presence of deletions, duplications, and chromosomal rearrangements at viral-host junctions (30). These genomic alterations strongly suggest that HPV integration is facilitated by error-prone repair mechanisms, with MMEJ as the most probable pathway. In contrast, HR is a high-fidelity repair pathway, and classical NHEJ typically preserves genomic sequences at breakpoints, making these pathways less likely as the predominant mechanism driving HPV integration. Further supporting this, mutational signature analysis of predominantly HPV-positive cervical cancers reveals a high frequency of ID6 (31). ID6 is the characteristic insertion/deletion signature associated with POLQ-mediated MMEJ, marked by microhomology usage at deletion junctions (31, 32). Another study also revealed that HPV-positive tumors exhibited a higher percentage of MMEJ repair scars (33). The distinct mutational signatures suggest that MMEJ activation in HPV-positive cancer is likely driven by viral oncogene rather than simply arising as a compensatory response to inherent HR deficiency.

Indeed, our findings indicate that HPV16 E6’s preferential use of MMEJ for integration results from its ability to modify the repair environment within HPV-infected cells. First, as our data demonstrates, E6 directly promotes MMEJ activity by upregulating POLQ. Second, E6 exacerbates this pre-existing bias towards MMEJ by inducing replication stress and consequent DSB accumulation that persists into mitosis. Recent studies have established MMEJ as the primary DSB repair pathway active during mitosis, when HR and NHEJ are largely suppressed (16–18). Consistent with this, our previous study demonstrated that accumulation of DNA damage in mitosis, driven by Cyclin D1, also results in a critical cellular reliance on POLQ-mediated MMEJ for repair (34). Taken together, these findings reinforce the idea that the HPV-driven repair pathway is not merely a consequence of genomic instability but rather an actively regulated process orchestrated by E6, thereby facilitating viral genome integration and promoting tumorigenesis.

Our study uncovers a novel mechanism by which HPV16 E6 enhances MMEJ activity. Specifically, E6 activates the E3 ligase UBE3A and redirects it towards the proteasomal shuttle protein RAD23A, consequently stabilizing POLQ. Notably, RAD23A knock-out phenocopied E6 overexpression in elevating both POLQ levels and MMEJ activity, confirming that RAD23A is a primary target through which E6 regulates MMEJ. The HPV E6 oncoprotein is known to function as an allosteric activator of the E3 ligase UBE3A, thereby altering its substrate profile. E6-induced degradation of p53 is the most well-characterized consequence of this interaction (10). In addition to p53, RAD23A has also been identified as another important UBE3A substrate, and previous studies have shown that E6 enhances UBE3A-dependent ubiquitination of RAD23A (22–24).

RAD23A acts as a proteasomal shuttle, using its UBA domains to bind ubiquitinated substrates and its UBL domain for proteasomal recruitment (27). A cornerstone of our mechanistic model is the RAD23A-mediated suppression of POLQ, as evidenced by experiments employing the proteasome inhibitor MG132. MG132 treatment not only stabilized POLQ but also markedly enhanced the interaction between RAD23A and POLQ, which implies that RAD23A preferentially associates with ubiquitinated POLQ, consistent with its role as a proteasomal shuttle factor.

RAD23A itself is also subject to regulation. While UBE3A-mediated ubiquitination has been linked to the degradation of RAD23A (23), recent evidence suggests that in addition to this role, it also acts as a functional regulator. Introduction of multiple arginine-to-lysine substitutions in the lysine desert region of RAD23A, within the UBL domain, promotes the ubiquitination of RAD23A but compromises RAD23A’s function (24). The differential effects of wild-type RAD23A and the ΔUBL mutant confirm that the UBL domain is indispensable in this process. While wild-type RAD23A overexpression suppressed both POLQ levels and MMEJ activity, the ΔUBL mutant, which is unable to recruit the proteasome, failed to do so, instead behaving as a dominant-negative factor, which enhanced MMEJ. This dominant-negative behavior provides critical evidence for our model. The ΔUBL mutant, while retaining its UBA domains for binding ubiquitinated substrates, lacks the essential UBL domain required for proteasomal interaction and efficient cargo handoff. By acting as a dominant-negative trap, the ΔUBL mutant effectively sequesters ubiquitinated POLQ from proteasomal degradation.

Despite elucidating a critical mechanism by which HPV16 E6 promotes MMEJ, this study has several limitations and leaves some questions unresolved. First, our model demonstrates that E6 activates UBE3A to prevent the degradation of ubiquitinated POLQ by compromising the RAD23A shuttle. However, the specific host E3 ubiquitin ligase that initially targets POLQ for ubiquitination remains unidentified. Second, due to the technical challenges of generating infectious HPV virions *in vitro*, the functional assessments relied on reporter-based systems for MMEJ activity and DNA integration efficiency. While informative, these systems may not fully recapitulate the complexities of authentic viral genome integration within the intricate host cell environment.

The identification of POLQ stabilization as a critical, E6-driven event essential for integration suggests that this pathway could be therapeutically exploited. Currently, prophylactic HPV vaccines are highly effective at preventing new infections, but they offer no therapeutic benefit for individuals with existing, persistent high-risk HPV infections. Our findings provide a rationale for developing novel secondary prevention strategies aimed at this large, at-risk population. For instance, pharmacological inhibitors of POLQ’s enzymatic activity could potentially be employed to suppress the MMEJ pathway. Such an approach might significantly reduce the frequency of new viral integration events, thereby blocking the pivotal step required for malignant transformation and offering a new strategy to prevent carcinogenesis in infected populations.

In conclusion, this study highlights HPV16 E6 as a key regulator of DNA repair, driving the error-prone MMEJ pathway to facilitate viral integration. By promoting the ubiquitination of RAD23A via UBE3A, E6 upregulates POLQ, creating a favorable environment in the host genome for MMEJ-mediated integration. These findings illustrate how viruses exploit the flexibility of host repair pathways, with broad implications for both virology and cancer biology. Future studies investigating the regulation of MMEJ and its interactions with other repair pathways could uncover new approaches to prevent viral integration and reduce the risk of HPV-associated cancers.

## Materials and Methods

### Cell culture

All cell lines were grown at 37 ℃ with 5% CO2. HEK293T (ATCC CRL-3216), U2OS (ATCC HTB-96), and CAL33 cells were cultured in DMEM medium (Gibco #11965092). RPE cells were cultured in DMEM/F12 medium (Gibco #10565042). FaDu (ATCC HTB-43) and UPCI-SCC-154 (ATCC CRL-3241) cells were cultured in MEM medium (Gibco #11095098). SCC9 cells were cultured in DMEM/F12 medium supplemented with 400 ng/ml hydrocortisone (Sigma Aldrich #H0888). All media were supplemented with 10% fetal bovine serum (Sigma Aldrich #F2442) and 100 U/mL penicillin-streptomycin (Gibco #15140163). CAL33 and SCC9 cells were gifts from Dr. Ravindra Uppaluri.

### Plasmids constructs

The HPV16 E6 and HPV16 E7 constructs were generated by cloning the cDNAs from pB-actin 16 E6 (Addgene #13713) and cmv 16 E7 (Addgene #13686) into the pCDH vector with an N-terminal HA tag, respectively. UBE3A constructs were cloned from pcDNA3.1+/C-(K)DYK-UBE3A (Genscript #OHu22798) into the pCDH vector with an N-terminal 3xFLAG tag. POLQ constructs were cloned from pcDNA3.1+/C-(K)DYK-POLQ (Genscript #OHu02809) into the pcDNA3 vector with an N-terminal 3xFLAG tag. RAD23A constructs were cloned from pcDNA3.1+/C-(K)DYK-RAD23A (Genscript # OHu15837) into the pCDH vector with an N-terminal V5 tag. Mutants of HPV16 E6, UBE3A, and POLQ were generated by site-directed mutagenesis using the plasmids mentioned above as the template. pcDNA3-HA-Ubiquitin plasmid was a gift from Dr. Nicholas W. Ashton.

### Lentivirus production and infection

Lentivirus was produced in HEK293T cells using the calcium phosphate transfection method (Takara bio #631312). Per each 10-cm plate, solution A was made with 12 µg transfer plasmid, 10 µg psPAX2 (Addgene #12260), 3 µg pMD2.G (Addgene #12259), 50 µL 2 M Calcium Solution, and 425 µL of H2O, and mixed with solution B (500 µL 2X HBS). The mixture was incubated for 5 minutes at room temperature and added dropwise to plates. The culture medium was changed after 18 hours, and the virus was harvested 48 hours post-transfection. Cells were transduced with the virus in the presence of 8 µg/mL polybrene, followed by puromycin or blasticidin selection for 72-96 hours.

### Plasmid Transfection

Plasmid transfection was performed using Lipofectamine LTX with PLUS reagents (Invitrogen #15338100). Cells were seeded at 60–70% confluency one day prior to transfection. Plasmid DNA and PLUS Reagent were diluted in Opti-MEM, mixed with the diluted LTX at a 1:1 ratio, incubated at room temperature for 5 minutes, and added dropwise. Medium was replaced after 18 hours, and cells were harvested 48 hours post-transfection.

### CRISPR/Cas9-mediated knockout

Gene knockout mediated by CRISPR/Cas9 was achieved by either lentiviral infection or electroporation of Cas9-sgRNA ribonucleoprotein (RNP) complex. For lentiviral infection, sgRNA sequence was cloned into the lentiCRISPR v2 (Addgene Plasmid #52961) via ligation (Takara Bio #6023) and packaged to make lentivirus. For electroporation, recombinant sgRNA (Integrated DNA Technologies), Alt-R® S.p. Cas9 Nuclease V3 (Integrated DNA Technologies, #1081059), and Enhancer (Integrated DNA Technologies, #1075915) were mixed and incubated at room temperature for 10 minutes to allow RNP formation. The RNP mixture was then added to the cell suspension, and electroporation was performed using the 4D-Nucleofector System (Lonza). For clonal selection, cells were subjected to limiting dilution to isolate single-cell clones. Single-cell clones were isolated by limiting dilution and validated by immuoblot analysis or Sanger sequencing. sgRNAs sequences used in this study are listed in Supplementary Methods.

### Western blot

Proteins were extracted by lysing cells in RIPA buffer (Cell Signaling Technology, #9806S) supplemented with protease and phosphatase inhibitor cocktails (Cell Signaling Technology, # 5872S) and PMSF (Cell Signaling Technology, #8553S). Proteins were denatured by boiling at 70 °C for 10 minutes in 2× Laemmli sample buffer (Santa Cruz, #sc-286962), separated on NuPAGE Tris-Acetate gels (Invitrogen #EA0375BOX) or Bis-Tris gels (Invitrogen #NP0321BOX), and transferred onto 0.45 μm nitrocellulose membranes (Biorad #1620115).

Membranes were blocked with 5% milk in 1X TBST (Tris-buffered saline [pH 7.4] with 0.1% Tween 20) for 1 hour at room temperature, incubated with primary antibodies overnight at 4 °C, washed, and incubated with secondary antibodies for 1 hour at room temperature. Membranes were imaged using Amersham Imager 600 following incubation with chemiluminescent substrate (Life Technologies #34580) or Li-Cor Odyssey CLx imaging system (LI-COR Biosciences). The information on antibodies is provided in the Supplementary Methods.

### Co-immunoprecipitation

Cells were lysed for 30 minutes in cell lysis buffer (Cell Signaling Technology, #9803) supplemented with Protease/Phosphatase Inhibitor Cocktail (Cell Signaling Technology, #5872) and MG-132 (Selleck #S2619). After three 30-s sonication pulses, lysates were centrifuged, and the supernatants were collected. Ten percent of each lysate was reserved as input, and the remainder was subjected to immunoprecipitation. Immunoprecipitations were performed with anti-FLAG M2 (Sigma-Aldrich, #F3165) or anti-V5 (Abcam, #ab27671) antibodies conjugated to Dynabeads Protein G (Invitrogen, #10004D). Beads were washed with lysis buffer, and bound proteins were eluted by boiling in 2× Laemmli sample buffer (Santa Cruz, #sc-286962) at 70 °C for 20 minutes.

### DNA Repair Reporter assay

Cells with integrated I-SceI endonuclease-based DSB repair reporter cassettes (MMEJ-GFP, DR-GFP, and EJ5 reporters) were infected with adenovirus expressing I-SceI, and medium was changed 5 hours post-infection. When indicated, the inhibitors were added when changing medium. 48 hours after infection, the activity of corresponding DSB repair pathway was determined as the proportion of GFP-positive cells by flow cytometry analysis on the CytoFLEX S flow cytometer (Beckman Coulter).

The CRISPR-based MMEJ repair cassette contains a GFP template with an insert containing a stop codon and a Cas9 recognition site flanked by a 7 bp microhomology sequence. Upon Cas9-induced cleavage, MMEJ-mediated repair restores GFP expression. Cells integrated with this reporter were infected by lentivirus expressing corresponding sgRNA. MMEJ activity was also determined as the proportion of GFP-positive cells by flow cytometry analysis 72 hours post-infection. Flow cytometry data were analyzed using FlowJo (v10.8, BD Biosciences).

### MMEJ-mediated double-stranded DNA integration reporter assay

In this system, we selected the C-terminus of the FBL gene as the integration target and transfected cells with two plasmids, including a donor plasmid containing a promoter-less GFP gene flanked by 20 bp homologous sequence and an sgRNA-expressing plasmid targeting both the FBL locus and the donor plasmid, enabling GFP integration into the FBL locus via MMEJ-mediated repair. GFP-positive cells were quantified by flow cytometry 48 hours post-transfection to determine the efficiency of MMEJ-mediated double-stranded DNA integration.

### Alkaline Comet assay

Cells were mixed with 50 L melted low-melting agarose and pipetted onto the CometSlides (R&D Systems #4250050K). Once the agarose solidified, the slides were immersed in lysis solution for 18 hours at 4 ℃, then incubated in alkaline unwinding solution for 1 hour to denature the DNA, and electrophoresed at at 21 V for 45 minutes in alkaline electrophoresis solution. The slides were stained with SYBR green solution, scanned with a fluorescence microscope (Zeiss Axio Observer), and analyzed using CometScore 2.0 software (TriTek Corp., Sumerduck, VA, USA).

### Immunofluorescence

Cells were fixed with 4% paraformaldehyde in PBS for 15 minutes at room temperature and permeabilized with 0.3% Triton X-100 in PBS for 10 minutes, followed by blocking with 3% milk in PBS for 1 hour at room temperature. The slides were incubated with anti-V5 antibodies (Cell Signaling Technology, #13202S) at 4℃ overnight, followed by incubation with secondary antibodies (Invitrogen, #A-11008) for 1 hour at room temperature. Nuclei were counterstained with NucBlue (Invitrogen, #P36981). The slides were scanned using a fluorescence microscope. At least 100 cells were counted for each sample. Foci quantification was performed using CellProfiler software (version 4.2.6).

### Cell cycle synchronization

Cells were incubated with 8 µM CDK1 inhibitor RO-3306 (Selleckchem #S7747) for 16 hours to induce G2 phase arrest. Following synchronization, cells were washed twice with PBS and released into fresh medium for 1 hour to allow progression into early mitosis. After release, cells were incubated with 100 ng/mL nocodazole (Sigma Aldrich #M1404) for 4 hours. Successful synchronization was confirmed by immunoblot analysis of cyclin B1.

### Clonogenic assay

Mitotically arrested U2OS cells were seeded into 6-well plates and treated with DMSO or 20μM ART558 (MedChemExpress, #HY-141520). 10 days after treatment, colonies were fixed with methanol and acetate (3:1 v/v) for 1 hour, followed by staining with crystal violet (0.5% w/v) for 1 hour. Colonies were imaged using the Amersham Imager 600 and quantified with ImageJ software (National Institutes of Health, Bethesda, MD, USA).

### Statistical analysis

Data analyses and visualization were performed using GraphPad Prism 10.2.2 (GraphPad Software, San Diego, CA, USA). Comparisons between two groups were conducted using unpaired two-tailed Student’s t-test, while multiple-group comparisons were analyzed using one-way ANOVA with Tukey’s post hoc test. Statistical significance was defined as p < 0.05.

## Acknowledgments and funding sources

We thank members of the D’Andrea laboratory for their helpful suggestions and comments. This work was supported by the Breast Cancer Research Foundation (A.D.D.), the Fanconi Anemia Research Fund (A.D.D.), the Ludwig Center at Harvard (A.D.D.), and the Smith Family Foundation (A.D.D.). This work was also supported by the SENSHIN Medical Research Foundation (S.A.), the Japan Society for the Promotion of Science Overseas Research Fellowships (S.A.).

