## Supplementary Information for "HPV16 E6 oncoprotein promotes microhomology-mediated viral integration by upregulating POLQ expression"

#### Supporting Information Text

#### Extended Methods.

##### sgRNA sequences

The sequences of single-guide RNAs (sgRNAs) used for CRISPR/Cas9-mediated gene knockout or targeting experiments are listed below.

| Target Gene | Target sgRNA sequence |
|---|---|
| ***UBE3A*-1** | AAACAAGAAAGGTCCTCGAG |
| ***UBE3A*-2** | CTAGCTAGAGATGATCGCTA |
| ***UBE3A*-3** | TGAGCTATCACCTATCCTTG |
| ***UBE3A*-4** | CTACTACCACCAGTTAACTG |
| ***TP53*** | CCATTGTTCAATATCGTCCG |
| ***POLQ*** | TCTGATCAATCGCCTCATAG |
| ***E6*-3** | AATATAAGGGGTCGGTGGAC |
| ***E6*-4** | GAAACATTGCAGTTCTCTTT |
| ***E7*-1** | ATTAAATGACAGCTCAGAGG |
| ***E7*-2** | AAGCGTAGAGTCACACTTGC |
| ***FBL*** | CTCTCACAGGCCACCCCCCA |

##### Antibodies Used for Immunoblot

A list of antibodies used for immunoblot, along with their sources, dilution ratio, and catalog numbers, is provided below.

| Immunoblot Antibodies | Source | Dilution | Identifier |
|---|---|---|---|
| HA | Cell Signaling Technology | 1:1000 | #3724S |
| V5 | Cell Signaling Technology | 1:1000 | #13202S |
| Flag | Sigma Aldrich | 1:1000 | #F1804 |
| GAPDH | Cell Signaling Technology | 1:1000 | #5174S |
| UBE3A | Cell Signaling Technology | 1:1000 | #7526S |
| p53 | Santa Cruz Biotechnology | 1:200 | #SC-126 |
| PARP1 | Abcam | 1:1000 | #ab32138 |
| LIG3 | Abcam | 1:1000 | #ab185815 |
| XRCC1 | Cell Signaling Technology | 1:1000 | #76998S |
| HMCES | Atlas Antibodies Ab | 1:1000 | #HPA044968 |
| APEX2 | Cell Signaling Technology | 1:1000 | #74728S |
| FEN1 | Cell Signaling Technology | 1:1000 | #82354S |
| $\gamma$–H2AX | Cell Signaling Technology | 1:1000 | #9718S |
| RPA | Cell Signaling Technology | 1:1000 | #52448S |
| Phosphor-RPA32/RPA2 (S8) | Cell Signaling Technology | 1:1000 | #83745S |
| Cyclin B1 | Cell Signaling Technology | 1:1000 | #4138S |
| POLQ | Cell Signaling Technology | 1:3000 | #64708 |
| RAD23A | Cell Signaling Technology | 1:1000 | #24555 |
| V5 | Abcam | 1:1000 | #ab27671 |
| HRP-coupled anti-mouse IgG antibody | Cell Signaling Technology | 1:10,000 | #7076V |
| HRP-coupled anti-rabbit IgG antibody | Cell Signaling Technology | 1:10,000 | #7074S |
| DyLight 680-coupled anti-mouse IgG antibody | Cell Signaling Technology | 1:10,000 | #5470S |
| DyLight 800-coupled anti-rabbit IgG antibody | Cell Signaling Technology | 1:10,000 | #5151S |

**Quantitative Real Time-PCR**

The whole RNA was extracted using the RNeasy Mini Kit (Qiagen # 74104) followed by reverse transcription with High-Capacity RNA-to-cDNA Kit (Applied Biosystems # 4387406). The following primers were used in an RT-PCR reaction using Power SYBR Green PCR MasterMix (Applied Biosystems # 4367659).

| qPCR Primers | Forward Primer Sequence | Reverse Primer Sequence |
|---|---|---|
| ***POLQ*** | ACCTCTCCATCAAGGCATTTCT | GCAAAAGTTCCAGCAGATACCC |
| ***GAPDH*** | GGAGCGAGATCCCTCCAAAAT | GGCTGTTGTCATACTTCTCATGG |

**Figures**

### Supplementary Figure 1

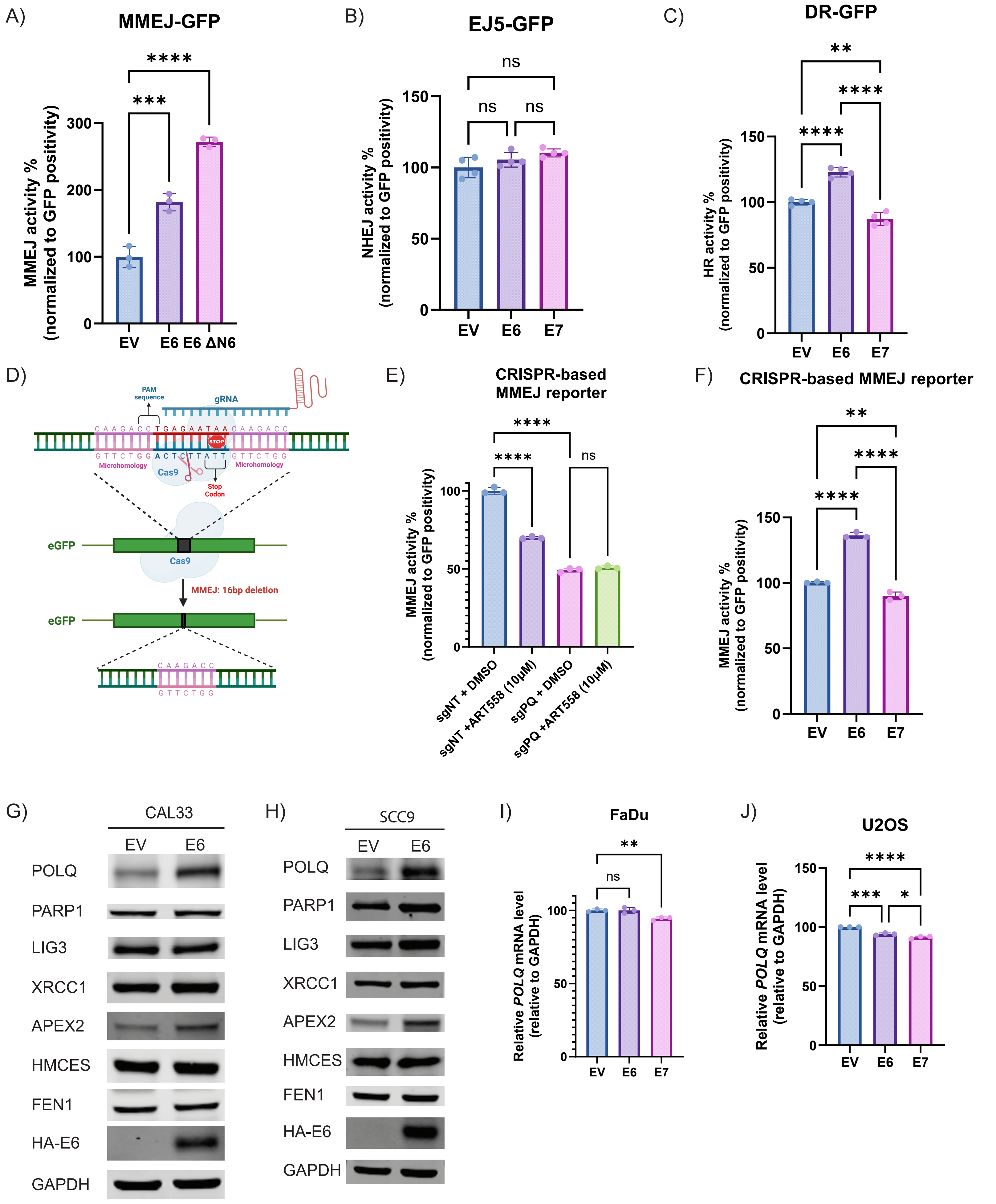

**Fig. S1.** HPV16 E6 specifically enhances MMEJ activity by increasing POLQ protein level.

A) Relative microhomology-mediated end joining (MMEJ) activity measured using the I-SceI-based MMEJ reporter assay in U2OS cells expressing empty vector, HPV16 E6, or HPV16 E6 ΔN6. Data are shown as the mean ± SD. Statistical significance was calculated using one-way ANOVA with Tukey's post hoc test (***, $P < 0.001$; ****, $P < 0.0001$).

B) Relative non-homologous end joining (NHEJ) activity measured using an EJ5-GFP reporter assay in U2OS cells expressing empty vector, HPV16 E6, or HPV16 E7.

C) Relative homologous recombination (HR) activity measured using an EJ5-GFP reporter assay in U2OS cells expressing empty vector, HPV16 E6, or HPV16 E7. Data are shown as the mean ± SD. Statistical significance was calculated using one-way ANOVA with Tukey's post hoc test (**, $P < 0.01$; ****, $P < 0.0001$).

D) Schematic illustration of the CRISPR-based MMEJ GFP reporter construct.

E) Relative MMEJ activity measured using the CRISPR-based reporter assay in U2OS cells with or without *POLQ* knock-out, treated with DMSO or 10 µM ART558. Data are shown as the mean ± SD. Statistical significance was calculated using one-way ANOVA with Tukey's post hoc test (***, $P < 0.001$; ****, $P < 0.0001$).

F) Relative MMEJ activity measured using the CRISPR-based reporter assay in FaDu cells expressing empty vector, HPV16 E6, or HPV16 E7. Data are shown as the mean ± SD. Statistical significance was calculated using one-way ANOVA with Tukey's post hoc test (**, $P < 0.01$; ****, $P < 0.0001$).

G-H) Immunoblot of POLQ, HA-tagged E6, other key MMEJ pathway components (PARP1, LIG3, XRCC1, APEX2, HMCES, FEN1) and GAPDH in HPV-negative cell lines, G) CAL33 and H) SCC9, expressing empty vector, HPV16 E6, or HPV16 E7.

I-J) Relative *POLQ* mRNA expression level (normalized to GAPDH) in measured by qRT-PCR in I) FaDu and J) U2OS cells expressing empty vector, HPV16 E6, or HPV16 E7. Data are shown as the mean ± SD. Statistical significance was calculated using one-way ANOVA with Tukey's post hoc test (*, $P < 0.05$; **, $P < 0.01$; ***, $P < 0.001$; ****, $P < 0.0001$).

#### Supplementary Figure 2

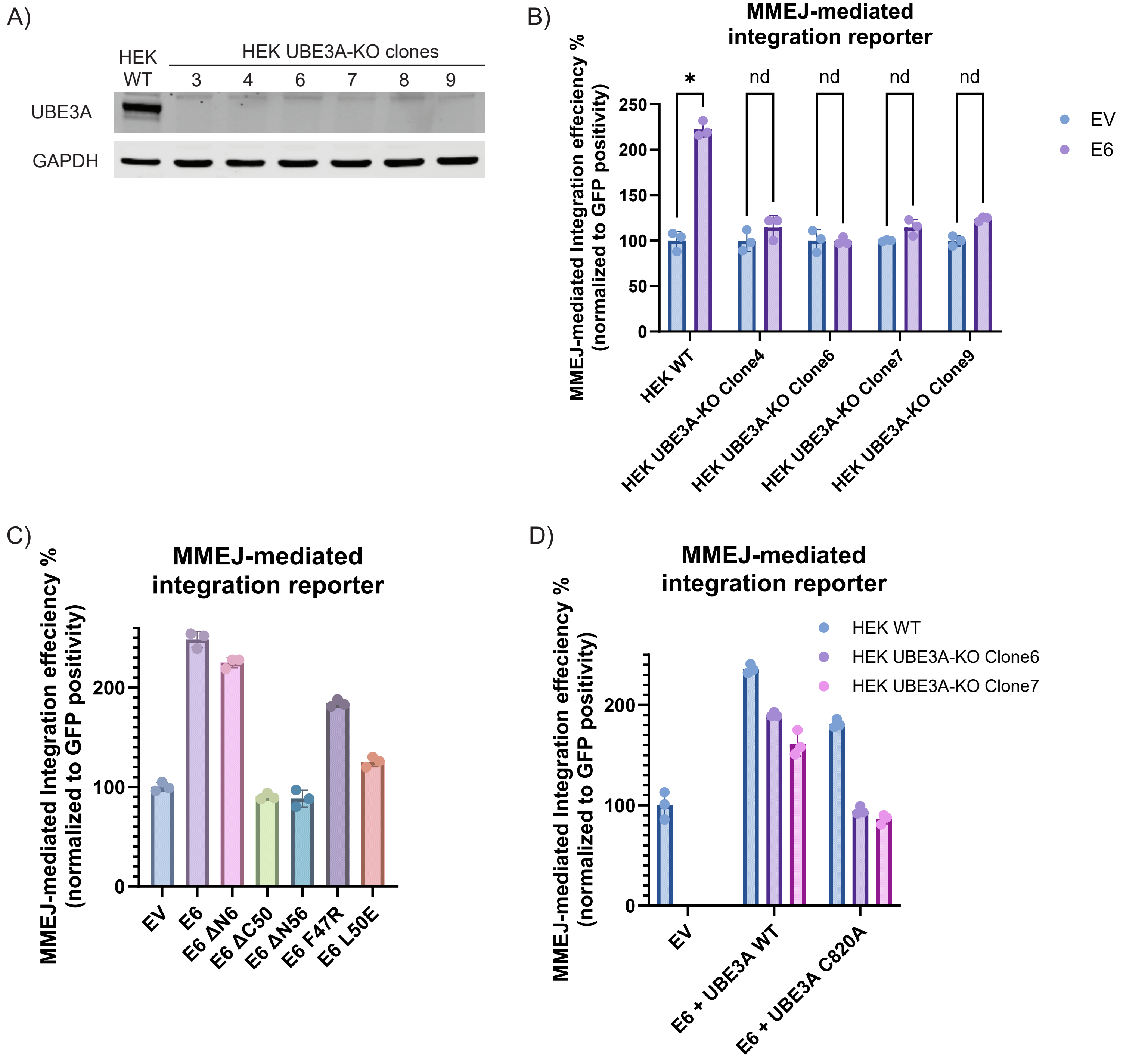

**Fig. S2.** HPV16 E6-mediated DNA integration is dependent on the E3 ligase activity of UBE3A.

A) Immunoblot of UBE3A and GAPDH in wild type HEK293T cells and different *UBE3A* knock-out HEK293T clones.

B) Relative microhomology-mediated end joining (MMEJ)-mediated integration efficiency measured using the MMEJ-mediated integration reporter in wild type HEK293T cells and different *UBE3A* knock-out HEK293T clones.

C) Relative MMEJ-mediated integration efficiency measured using the MMEJ-mediated integration reporter in wild type HEK293T cells expressing EV, E6, wild-type E6ΔN6 and its mutants. Data are shown as the mean ± SD. Statistical significance was calculated using two-tailed Student's t-test (*, $P < 0.05$; nd, no difference).

D) Relative MMEJ-mediated integration efficiency measured using the MMEJ-mediated integration reporter in wild type HEK293T cells and different *UBE3A* knock-out HEK293T clones expressing empty vector, wild-type UBE3A and catalytically inactive UBE3A (C820A).

#### Supplementary Figure 3

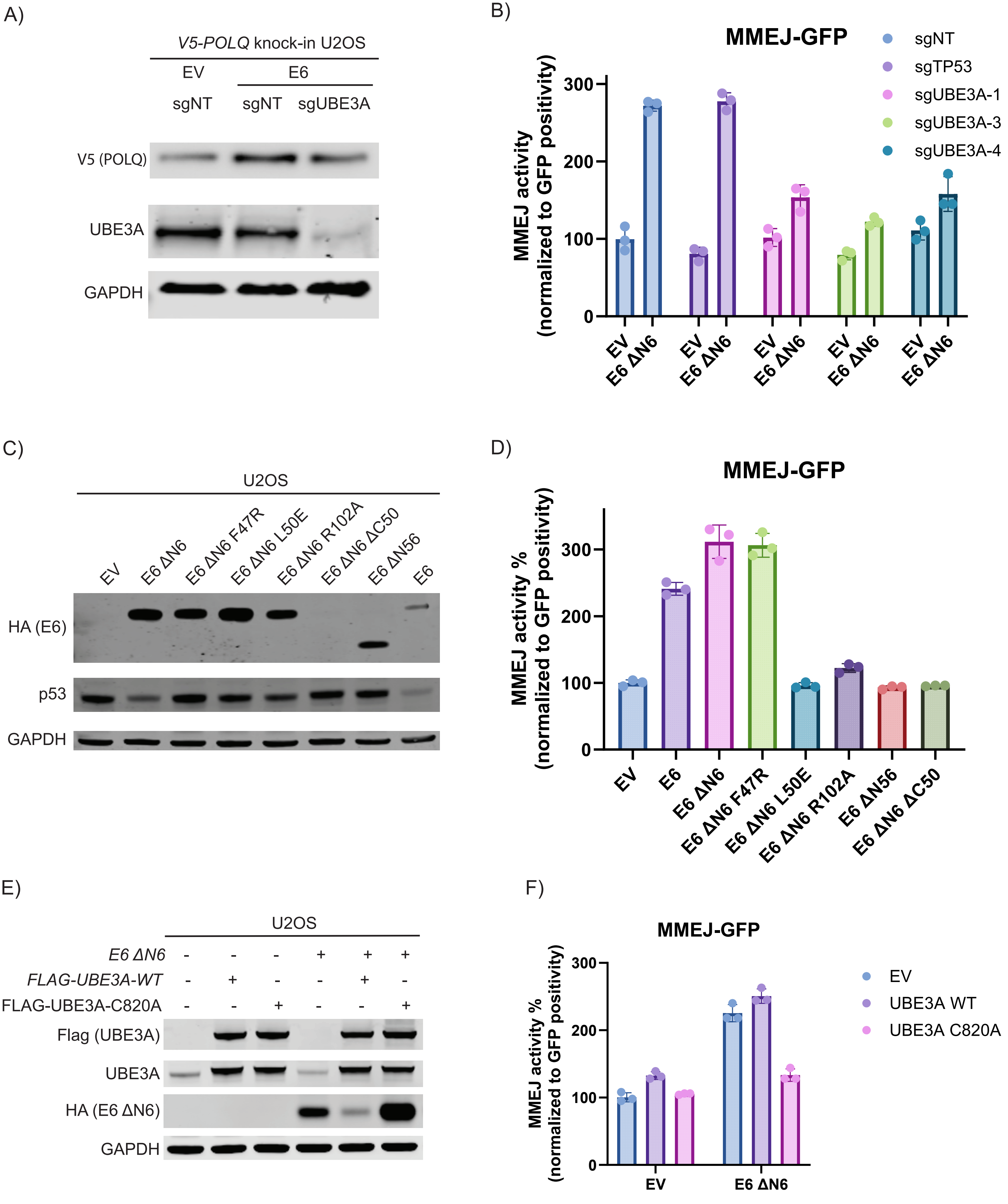

**Fig. S3.** HPV16 E6 increases MMEJ activity and POLQ level in a UBE3A-dependent manner.

A) Immunoblot of endogenous POLQ, HA-tagged HPV16 E6, UBE3A and GAPDH proteins in wild-type or UBE3A knock-out *V5-POLQ* knock-in U2OS cells expressing empty vector or wild-type HPV16 E6.

B) Relative microhomology-mediated end joining (MMEJ) activity measured using the I-SceI-based MMEJ reporter assay in wild-type or *UBE3A* knock-out U2OS cells expressing empty vector or HPV16 E6 ΔN6.

C) Immunoblot of HA-tagged E6 ΔN6 and its mutants, p53 and GAPDH proteins in wild-type U2OS cells expressing empty vector, wild-type HPV16 E6 ΔN6, and the indicated E6ΔN6 mutants.

D) Relative MMEJ activity measured using the I-SceI-based MMEJ reporter assay in U2OS cells expressing empty vector, wild-type HPV16 E6, wild-type HPV16 E6 ΔN6, and the indicated E6ΔN6 mutants.

E) Immunoblot of FLAG-tagged wild-type or catalytically inactive (C820A) UBE3A, HA-tagged E6, and GAPDH in U2OS cells expressing empty vector, wild-type or C820A mutant UBE3A, with or without HPV16 E6 ΔN6 expression.

F) Relative MMEJ activity measured using the I-SceI-based MMEJ GFP reporter assay in U2OS cells expressing empty vector, wild-type UBE3A or C820A mutant, with or without HPV16 E6 ΔN6 expression.

### Supplementary Figure 4

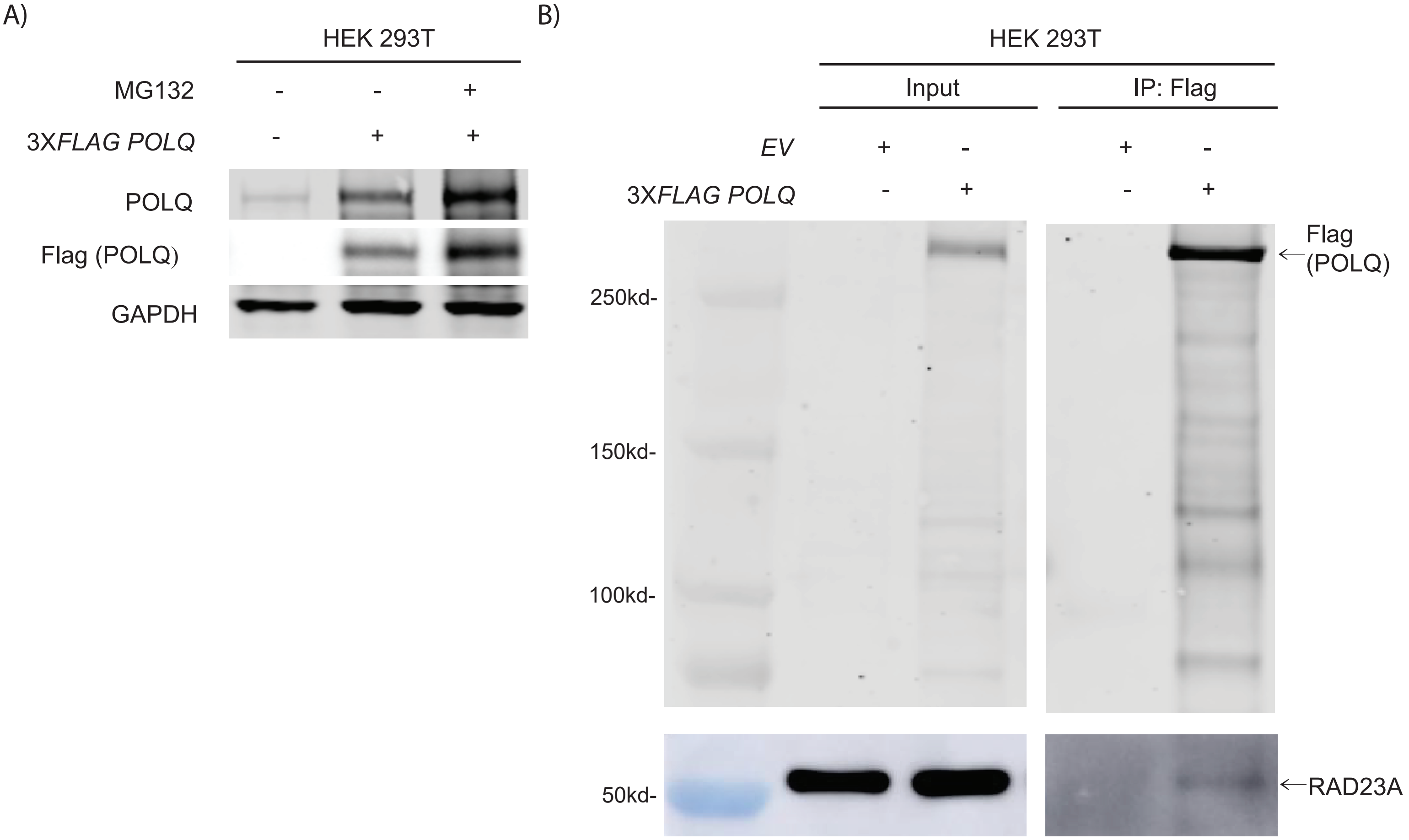

**Fig. S4.** Validation of POLQ stabilization and its interaction with endogenous RAD23A.

A) Immunoblot of FLAG-tagged POLQ in HEK293T with or without 3x*FLAG-POLQ* transfection and with or without MG132 treatment.

B) Co-immunoprecipitation analysis. Whole cell lysates from *POLQ* knock-out HEK293T cells reconstituted with empty vector or 3x*FLAG*-tagged full-length *POLQ*, were immunoprecipitated with anti-Flag antibody, followed by Immunoblot of input and IP samples for Flag (POLQ fragments) and RAD23A.
